# C-Terminal Amidation: Structural Insights into Enhanced Antimicrobial Peptide Efficacy and Amyloidogenesis

**DOI:** 10.1101/2025.03.24.645029

**Authors:** Amit Chaudhary, Anup K. Prasad, Lisandra L. Martin, Ajay Singh Panwar

## Abstract

Post-translational modifications, such as C-terminal amidation, are energetically costly but critical for membrane-active antimicrobial peptides (AMPs). This study examined the impact of C-terminal amidation on uperin 3.5 (U3.5, an amyloid-forming AMP) interactions with POPE:DOPG (3:1) lipid bilayers using molecular simulations. Whereas, monomeric U3.5-NH_2_ rapidly attached to the lipid bilayer surface, forming a stable α-helix, U3.5-OH exhibited weaker interactions. Simulations of U3.5 tetramers, derived from the amyloid cryo-electron microscopy structure, revealed that amidation enhanced peptide-bilayer and peptide-peptide interactions, initially stabilising β-sheet structures and facilitating embedding into the bilayer. The β-sheet tetramer gradually dissociated into monomeric U3.5-NH_2_ peptides on the bilayer surface. Following dissociation, the monomers formed stable, amphipathic α-helices that were strongly embedded in the bilayer, consistent with a carpet-like antimicrobial mechanism. Enhanced peptide-lipid interactions resulted in lipid redistribution and changes to membrane curvature, potentially leading to membrane rupture. These findings highlight the role of U3.5-NH_2_ amyloid as a “carrier vehicle” for antimicrobial action at the bilayer interface, emphasizing the crucial function of C-terminal amidation in stabilising peptide structure and promoting antimicrobial activity.

## Introduction

Antimicrobial peptides (AMPs) are expressed across most forms of life, including animal, plant, fungi, and bacteria, providing a first line of defense towards invading organisms^1^. AMPs are extremely diverse in terms of amino acid composition, length, secondary structures, activities, and selectivity, and although not exclusive, the majority are linear, cationic peptides, typically 10-50 amino acids long, with a significant proportion of hydrophobic residues resulting in the capacity to form amphipathic α-helices if in contact with lipid environments. Mechanistically, the largest class of AMPs typically disrupt bacterial membranes via a carpet (detergent-like) mechanism or insertion into the lipid bilayer to form pores (barrel-stave or toroidal pores) with deleterious effects on the bacteria^2,3^. As these AMPs target the bacterial membrane, which in contrast to eukaryotic species are anionic, AMPs offer enormous potential as future armory against multi-drug resistant bacteria^4^.

During an exploration of the AMP activity of glandular secretions from the Australian toadlet *Uperoleia mjobergii*, 20 AMPs were isolated and found to display antibiotic activity. 4 Of these, four peptides which belong to the uperin 3 family have since been studied by us^5–12^ and others ^13–18^, identifying both antimicrobial activity and, in addition, a capacity to self-assemble into amyloid. These peptides are 17 amino acids long and are cationic with an overall 3+ charge. Interestingly, the X-ray structure of uperin 3.5 amyloid was first resolved for a C-terminal hydroxylated version (U3.5-OH), revealing a cross-α-amyloid structure^17^. Later, using cryo-electron microscopy (cryo-EM), the naturally secreted and C-terminal amidated wild-type peptide (U3.5-NH_2_) was found to only form β-sheet rich polymorphs^16^. All members of the uperin 3 family of AMPs are naturally C-terminal amidated^18,19^. An enormous amount of energy is expended by an organism with each step of peptide biosynthesis as well as any post-translational modification of the peptide. Thus, the question arises as to the importance of this post-translational modification on the activity of the AMP, especially towards bacterial membranes.

Amidation is just one of the many post-translational modifications (PTM) observed in AMPs,^20^ and is the most common PTM at the C-termini. It has the function of reducing the overall peptide charge and in some cases, amidation is reported to be essential for membrane activity, typically associated with AMPs^21^. Intuitively, the structural impact of C-terminal amidation is unlikely to influence the overall propensity to form secondary structure elements compared with the same peptide containing a C-terminal carboxyl moiety. However, the amidation decreases the peptide charge (by 1+), suggesting an obvious impact on the electrostatic attraction towards the negatively charged bacterial membranes. The influence of the C-terminal amidation has attracted a number of studies with AMPs, concluding that C-terminal amidation assists to maintain a stable α-helix, maintaining an optimal angle for membrane penetration in model mammalian lipids^22^. Also, the carboxylated C-terminus leads to increased hydrogen bonding with the lipid phosphate head groups that maintain the peptides at the water-lipid interface, rather than penetrating the lipid bilayer^23^. In a further study of a short cationic AMP (Aurein 1.2), the authors examined the influence of C-terminal modification, i.e., Au-COOH, Au-CONH2, Au-CONH(CH3), on the membrane binding characteristics^24^. They concluded that the primary amide was important for peptide binding to the membrane, and this was found to be less affective for the other two modifications, attributing this to the hydration state of the amidation moiety.

Unlike these earlier studies on the role of amidation on the C-termini of AMPs, the U3.5 peptide can also self-assemble into amyloid, thus provides an additional opportunity to investigate the role of aggregated and disaggregation of AMP action at an interfacial bacterial-mimetic membrane bilayer. Studying the amyloid state interaction with the membrane rather than a single peptide is significant because amyloid assemblies exhibit distinct structural and functional properties that significantly influence membrane interactions. Unlike monomeric peptides, amyloid oligomers and fibrils can form higher-order structures that alter their binding affinity, insertion dynamics, and membrane-disrupting mechanisms. Enhanced membrane disruption has been observed for amyloid aggregates that can induce greater membrane destabilisation than monomers either due to their ability to form pore-like structures or disrupt lipid packing over a larger surface area^25^. Increased stability and persistence of amyloid states which are typically more stable and resistant to degradation, potentially leading to prolonged interactions with bacterial membranes, which could enhance antimicrobial efficacy^26^. Cooperative effects of amyloid-like assemblies can also exhibit synergistic effects, where multiple peptides act together to achieve stronger membrane perturbation than individual peptides^27^. Similarities and relevance to natural antimicrobial mechanisms are found for example, many host-defence peptides and amyloidogenic antimicrobial peptides leverage self-assembly to enhance their biological function, suggesting that the amyloid state plays a key role in their activity^28^. Finally, there are resistance considerations of amyloid in which bacteria may develop resistance to single peptides more easily than to higher-order peptide assemblies, which rely on multivalent interactions and mechanical disruption rather than mechanisms including specific receptor binding^29^. Thus, by studying amyloid state interactions with membranes, we can gain a more comprehensive understanding of peptide-membrane dynamics, leading to improved design strategies for antimicrobial therapies.

The amphibian U3.5 wt is an antimicrobial peptide consisting of 17-amino acids (GVGDLIRKAVSVIKNIV-NH_2_) that is secreted on the skin of the toadlet *Uperoleia mjobergii*. This cationic peptide has broad-spectrum antimicrobial activity against gram-positive bacteria. Previous experimental studies have shown that U3.5 exists in three different structures under different environmental conditions: a random coil in solution^7^, an α-helical amyloid fibril in the presence of lipids,^17^ and a β-sheet rich amyloid fibril in the absence of lipids^5–10,30,31^. Here, we employ fully atomistic molecular dynamics (FA MD) simulations to examine C-termini amidated and non-amidated U3.5 in monomeric and aggregated (tetrameric) states near a membrane bilayer composed of lipids in a ratio that simulates a negatively charged bacterial membrane.

### Simulation Method

The FA MD simulations were established using a anionic lipid bilayer consists of POPE:DOPG 3:1 with identical U3.5 peptide sequences containing either amidated or non-amidated C-termini, designated as U3.5-NH_2_ (or U3.5AM) and U3.5-OH (or U3.5NA), respectively. In these simulations, either a single (unstructured monomer) peptide or a tetrameric (β-sheet) amyloid of each peptide type was placed close and parallel to the lipid bilayer surface, and structural changes were monitored during these simulations, as shown in **Figure S1**. The orientation of the peptide was horizontal to the membrane. For the current study, we generated the random coil structure of a single U3.5 from a previous simulation^5^ and extracted parallel tetramer structure of U3.5 from the cryo-EM 10-layered cross-beta fibril structure^16^.

#### Selection and construction of membrane systems

The effect of C-terminal amidation on U3.5 peptide (U3.5-NH_2_ *versus* U3.5-OH) on the interaction with a model gram-negative bacterial membrane (composed of POPE and DOPG) was investigated using MD simulations (**Figure S1**). Different systems comprising either amidated or non-amidated U3.5 peptides and a model lipid bilayer were simulated (**Table S1**). Both peptide monomers and amyloid-derived peptide tetramers were considered. A lipid mixture corresponding to a 3:1 ratio of POPE (1-palmitoyl-2-oleoyl phosphatidylethanolamine) and DOPG (1-palmitoyl-2-oleoyl-*sn*-glycero-3-(phospho-rac-(1-glycerol))) was chosen to mimic a model gram-negative bacterial membrane (overall negatively charged)^32–34^. The Membrane Builder module of the CHARMM-GUI ^35,36^ server was utilized for the preparation of peptide-membrane systems. The membrane normal was assumed to be parallel to the *Z*-axis, and its center was located at *Z* = 0. Peptide monomers and tetramers were placed at a vertical separation of 15 Å and 20 Å, respectively, from the membrane surface. The input geometry for the heterogeneous membrane system was specified with lipid moieties as per the membrane system ratio in both upper and lower leaflets. The electrical neutrality of the system was maintained by the addition of appropriate numbers of sodium (Na^+^) or chloride ions (Cl^−^). The membrane-peptide system was solvated with TIP3P water molecules, to which Na^+^ and Cl^-^ ions were further added to achieve an ionic strength of 0.15 M NaCl. Simulation box volumes and the total number of atoms for individual systems are mentioned in **Tables S1** and **S2** of the Supplementary Information. The simulations were carried out under NPT conditions at 1 bar, and 310 K. Periodic boundary conditions were applied along all directions, and the Particle Mesh Ewald method was used for evaluating long-range electrostatic interactions. A constant pressure was maintained by using semi-isotropic coupling (in the *X* and *Y* directions) using the Berendsen barostat ^37^ at 1 bar. The system temperature was maintained at 310 K in all model membranes (**Table S2)**. All simulations were performed using the CHARMM 36m (C36m) all-atom force field^38,39^. An integration time step of 2 fs was used. VMD was used to visualize trajectories, and in-house scripts were used for the analysis of secondary structure, peptide distance from lipid bilayer, RDF plot, hydrogen bond, and energy plot.

## Results and Discussion

Approximately, 20% of all naturally occurring AMPs listed on the APD website (https://aps.unmc.edu), are C-termini amidated^40^. The Uperin 3 family of AMPs are all naturally C-terminal amidated ^10,13,19,41^, and to varying extents self-assemble into amyloid in the presence of salts^5–8,10,12,31,41^, although in pure water the peptides are unstructured. Of the Uperin 3 peptides, U3.5 has been the most intensively investigated ^5,7–10,12,16,17,31,41^, in a range of media and using a wide range of bioanalytical and computational methods.

### Monomer simulations

Snapshots from the simulation trajectory of the monomeric U3.5-NH_2_ peptide showed rapid attachment to the lipid surface with the formation of an amphipathic α-helix lying with the hydrophobic interface oriented towards the lipid head groups within 500 ns (**Figure 1a**). In another 10-20 ns, the U3.5-NH_2_ monomer ‘tumbled over’, orienting the hydrophobic residues of the peptide toward the “oily” interior of the bilayer. This peptide orientation persisted for the remainder of the 1200 ns simulation, with only a slight further insertion into the lipid bilayer. In contrast, although the monomeric U3.5-OH (**Figure 1(b)**) bound rapidly to the lipid bilayer via the N-termini, only partial α-helices (3_10_- and short α-helical turns) were observed over a similar simulation timescale with no clear secondary structure or peptide insertion visible at 1200 ns (**Figure S2**). The distances of both the N- and C-termini from the lipid bilayer for each peptide are shown in Figure 1(c, d), illustrating the preferential location of the N-termini of both peptides towards the lipid. The plots clearly indicate the transition of the U3.5-NH_2_ peptide into a conformation that resulted in membrane association (≈500 ns in **Figure 1(c)**). In contrast, the C-terminus of U3.5-OH remained unattached to the bilayer and was fluctuating through the entire 1200 ns simulation (**Figure S3**). The orientation of the peptides with respect to the bilayer was described by the angle, *θ*, formed between the peptide’s principal vector (C_α_1 to C_α_17) and the normal vector of the membrane (**Figure 1(e)**). According to this definition (see **Figure S3(b)**), an angle close to 90° (observed for U3.5-NH_2_) signifies a peptide lying flat on the bilayer. Whereas, U3.5-NH_2_ lay flat on the bilayer (consistent with results of Figures 1(a) and 1(c)), large fluctuations in the angle for U3.5-OH indicated loose attachment with the bilayer.

**Figure 1:**
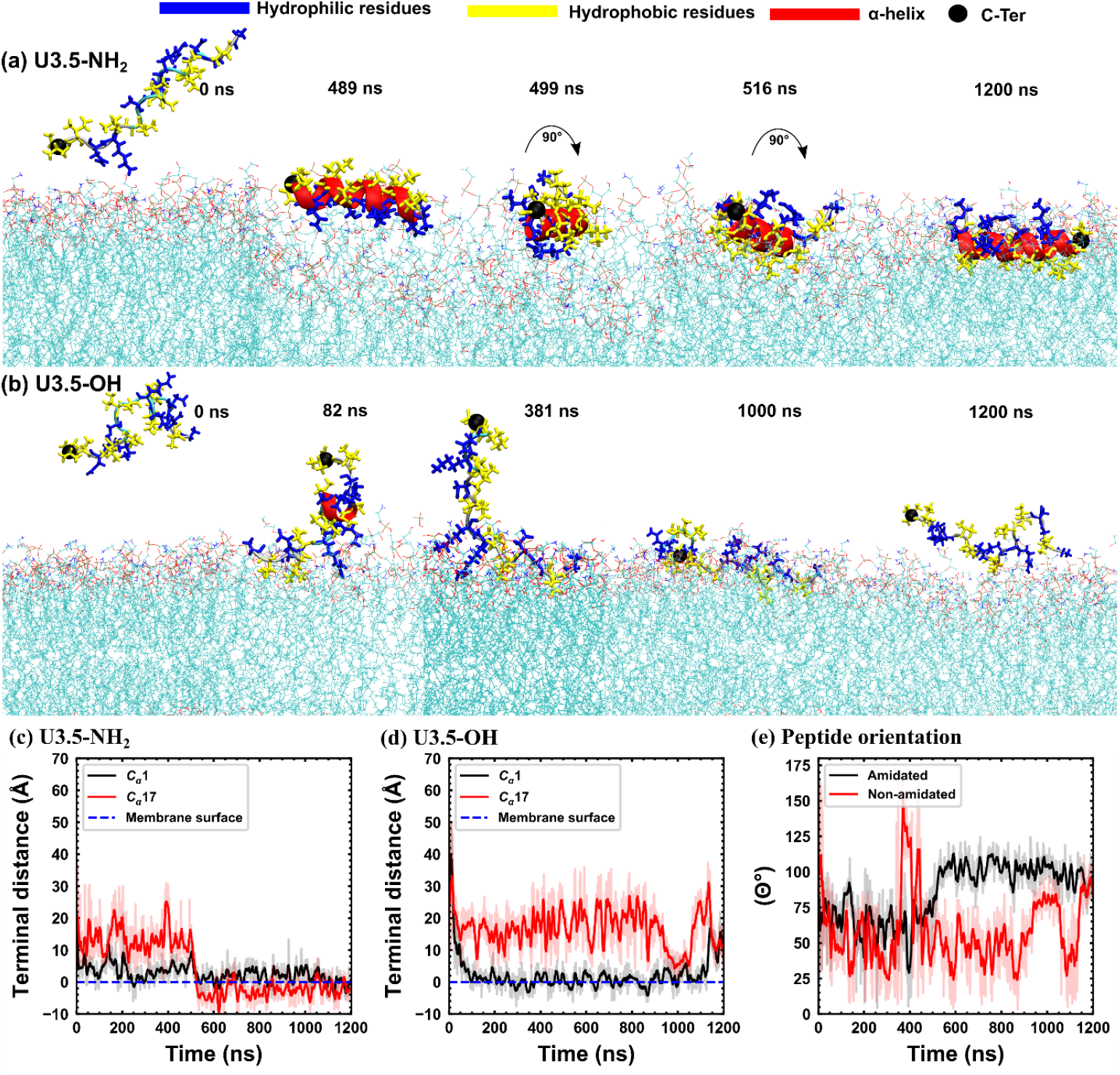
Secondary structure evolution of peptides at lipid bilayer during monomer simulations. (a) Snapshots of the U3.5-NH2 peptide depicting its adsorption onto the membrane surface and subsequent helical transition, initiated via interactions through the hydrophilic peptide face over 489 ns. The peptide undergoes a 180° rotation, enabling insertion of the hydrophobic peptide face into the lipid bilayer. (b) The U3.5-OH peptide similarly adsorbs onto the surface of the lipid bilayer; however, the negative charge at its C-terminus induces electrostatic repulsion, preventing interaction with the membrane or conformational transitions of the peptide. (c, d) Quantification of the distances of the peptides’ terminal alpha carbons (C_α_1 and C_α_17) from the membrane surface. (e) The orientation of the peptides, described by the angle formed between the peptide’s principal vector (C_α_1 to C_α_17) and the normal vector of the membrane is illustrated over the simulation time.

Additional insight into the approach and subsequent interactions of the two peptides with the membrane were obtained by examining changes in the average distance of the U3.5 peptide backbone (**C**_α_) atoms from the membrane surface throughout the simulation (**Figure S3**). Averaged **C**_α_ distances below < 5Å for the U3.5-NH2 peptide indicated strong and uniform interactions across the entire peptide length, signifying effective membrane association or binding. In contrast, the averaged distances for U3.5-OH were much larger, with larger deviations from the mean values, and increased with time from approximately 5Å to 15Å. These trends highlight the weaker overall membrane interaction for U3.5-OH. The U3.5-OH peptide primarily interacted via its N-terminus, particularly from C_α_1 to C_α_8, due to the presence of positively charged residues (ARG7 and LYS8) in this region. The negatively charged C-terminus, in the U3.5-OH case, was repelled from the membrane, resulting in limited membrane interaction. Two-dimensional electrostatic potential maps of the peptides positioned across the membrane (Z-axis) and the membrane surface (X-axis) further support increase in potential for U3.5-NH_2_ (**Figure S4**). The electrostatic potential map of the U3.5-NH_2_ peptide in the X-Z plane reveals a distinct positive region at X, Z = (−30, 20), corresponding to the location of the monomeric U3.5-NH_2_ (Figure S4(a)). Additionally, the potential profile along the Z-axis shows a clear difference in potential between the peptide center and the membrane, particularly near Z = 20. In contrast, the potential map for the U3.5-OH peptide-membrane (Figure S4(b)) lacks any clear distinction in between the electrostatic potential for the peptide centre and the membrane alone. This observation suggests a dilution of the electrostatic potential, likely caused by the C-terminus of the U3.5-OH peptide well above the membrane surface, thereby leading to weaker interaction with the lipid bilayer.

#### Tetramer simulations

Cryo-electron microscopy has revealed several polymorphs of the native U3.5-NH_2_ peptide^16^. One of these was a 3-blade propeller (with a 3-fold symmetry) of nine peptides with tight β-sheet interfaces^16^. Four adjacent U3.5-NH_2_ peptides from one of the “blades” of this structure were used as a β-sheet tetramer to probe the interaction of amyloid with the membrane bilayer. These simulations were run for 5 μs, significantly longer than for those used in monomer simulations. **Figures 2(a) and 2(b)** present snapshots comparing the secondary structure transitions of amidated and non-amidated β-sheet tetramers of U3.5, respectively. Interestingly, the U3.5-NH_2_ snapshots highlighted the initial tetramer binding parallel to the bilayer plane with both N- and C-termini in close contact with the lipid head groups until ∼ 4.5 μs, when one of the peptides began to dissociate or dissolve (**Figure 2(a)**). In contrast, the U3.5-OH peptide attached to the lipid bilayer through the N-termini with the C-termini (containing the carboxyl groups) oriented away for the membrane (**Figure 2(b)**). This was confirmed by comparing the average C-terminal distance of the respective tetramers from the membrane surface (represented by the dashed blue line in **Figure 2(c)**). The C-terminal distance showed a continuous decrease for U3.5-NH_2_ and turned negative beyond 4.5 μs, indicating the embedding of the peptide C-termini into the bilayer. For the U3.5-OH tetramer, the distance remained nearly uniform, supporting the observation of non-adsorbing C-termini in the tetramer snapshots in **Figure 2(b)**. Additionally, the analysis of tetramer orientations relative to the bilayer normal (as defined in Figure 1(e)) confirmed that while the U3.5-NH_2_ remained parallel to the bilayer surface, the U3.5-OH tetramer oriented away from the bilayer (**Figure 2(d)**). The evolution of β-sheet content in the tetramers over time exhibited distinct differences (**Figure 2(e)**). Whereas a sudden drop in -sheet content was observed for U3.5-OH tetramer within 500 ns, the β-sheet content of U3.5-NH2 tetramer remained largely stable until 4.0 µs when it precipitously declined. The β-sheet content in a tetramer is a direct measure of the number of inter-peptide hydrogen bonds in the tetramer. In the current context of C-terminus amidation, the β-sheet content is an indication of the extent of peptide-peptide interactions in the presence of a bilayer. As shown in **Figures 2(a)-2(d)**, C-terminus amidation influences the strength of interactions between the peptide tetramers and the bilayer. Whereas, C-terminus amidation resulted in strong adsorption and embedding of peptides into the bilayer, its absence resulted in a weakly adsorbed tetramer. Consequently, **Figure 2(e)** suggests that C-terminus amidation may influence the relative magnitudes of peptide-peptide (H-bond) and peptide-bilayer interactions. This interplay should have a direct impact on the secondary structure composition, particularly the β-sheet content of the U3.5 tetramer.

**Figure 2:**
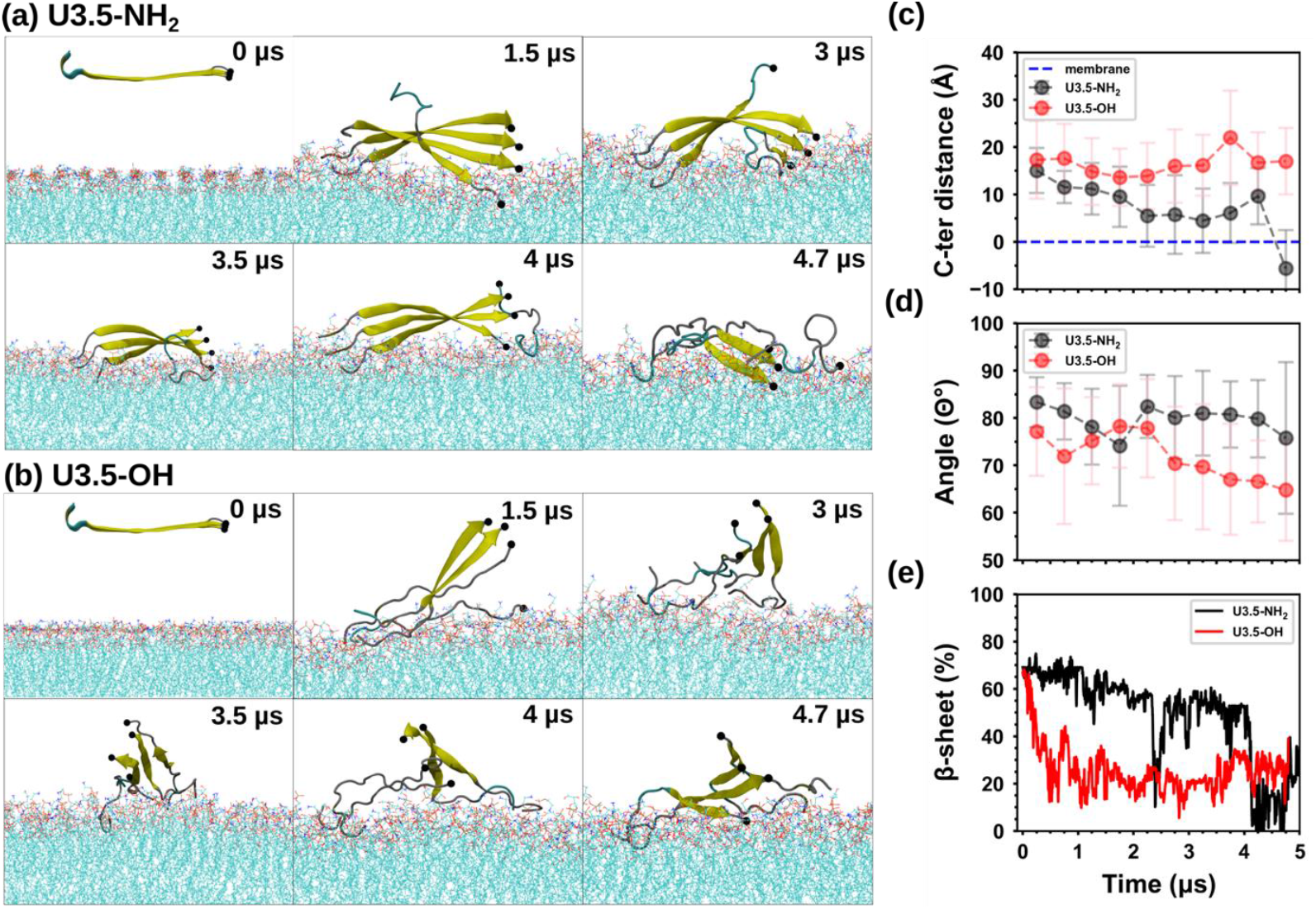
Comparison of secondary structure profiles between amidated and non-amidated U3.5 peptides in an initial β-sheet tetramer. Snapshots in both (a) and (b) highlight the transition states of these peptides, emphasizing their respective conformational changes and structural alterations. (c) illustrates the C-terminus distance for U3.5-NH_2_ and U3.5-OH peptide tetramers from the membrane surface (dashed blue color) over the simulation timescale. (d) This plot illustrates the orientation of peptides on the membrane surface over the simulation time, where 90º is normal to the lipid bilayer and 0º is parallel to the surface. (e) The average percentage of β-sheet structure present in the peptide tetramers over the simulation time (ns).

**Figure S5** shows the radial distribution functions (RDFs) corresponding to the distribution of the *C*_α_ atoms of specific residues (in the tetramer) relative to the phosphorus (P) atoms of the lipid headgroups. These were calculated for both U3.5-NH_2_ and U3.5-OH tetramers. The positively charged residues near the N-terminus - GLY1, ARG7, and LYS8 - exhibited strong and comparable association with the bilayer in both tetramers (**Figure S5(a)-(b)**). However, near the C-terminus, residues (LYS14 and VAL17) in the U3.5-NH_2_ tetramer showed stronger binding to the bilayer compared to their counterparts in the U3.5-OH tetramer (**Figure S5(c)-(d)**). This resulted directly from the electrostatic repulsion between the negatively charged C-termini of U3.5-OH and bilayer phosphate groups, weakening peptide-bilayer interactions. Plots of van der Waals (vdW) and electrostatic energy contributions for peptide-peptide and peptide-lipid bilayer interactions also highlight differences between U3.5-NH_2_ and U3.5-OH tetramers (**Figure S6**). Specifically, the peptide-peptide vdW energies were lower for the U3.5-NH_2_ tetramer, consistent with stronger inter-peptide H-bonds and more stable β-sheets in the amidated tetramer.

#### Extended simulations of dissociated U3.5-NH_2_ peptide from the tetramer

The gradual dissolution of the U3.5-NH_2_ tetramer started at ≈ 4.5 μs (shown in **Figure 2(a)**) with the separation of one peptide from the tetramer. Although, the simulation was extended to approximately 5.4 μs, the separated peptide remained in close proximity (8 Å) to the aggregate. Additionally, its secondary structure was predominantly random coil with negligible α-helix content (< 2%). This behaviour sharply contrasted with that of a single U3.5-NH_2_ peptide, which rapidly adopted an α-helical structure (≈40% within 200 ns, **Figure S2**) upon adsorption onto the lipid bilayer. The slow dynamics of secondary structure change for the detached peptide in the U3.5-NH_2_ tetramer was attributed to the strong peptide-peptide interactions on the bilayer surface. Given the behavior of a single U3.5-NH_2_ peptide and the gradual increase in the “peptide-trimer” separation distance, the detached peptide was expected to drift further from the trimer and ultimately adopt an α-helical conformation. This assumption was reasonable, as peptide-lipid electrostatic interactions were significantly stronger compared to peptide-peptide electrostatic interactions (**Figure S6(b) & (d)**). Stronger peptide-lipid interactions were expected to drive the dissolution of the U3.5-NH2 tetramer, leading to the embedding of amphipathic U3.5-NH2 peptides into the bilayer, similar to the behavior observed in **Figure 1**.

Therefore, to accelerate the dissolution and embedding process, the “peptide-trimer” separation distance was increased from 8 Å to 18 Å at 5.4 μs in the simulation shown in **Figure 2(a)**. A new 500 ns simulation was then initiated, with the “shifted” peptide defining the initial state. This process is described in Figure 3(a) through a series of snapshots representing: (i) end-state at 5.4 μs in the original U3.5-NH_2_ tetramer simulation, (ii) “artificial shifting” of the separated peptide by 10 Å, (iii) an intermediate snapshot at 350 ns showing the emergence of a 3_10_ helix in the separated peptide, and (iv) a snapshot at 500 ns showing the formation of an α-helix in the separated peptide. Significant secondary structure changes were observed in the dissociated peptide during the 500 ns simulation. In the first 200 ns, the single U3.5-NH_2_ peptide developed 3_10_-helices at the N-terminus (**Figures 3(b)** and **(c)**). These 3_10_-helices were precursors to α-helices that also emerged at the N-terminus after approximately 200 ns (**Figures 3(b)** and **(d)**).

**Figure 3:**
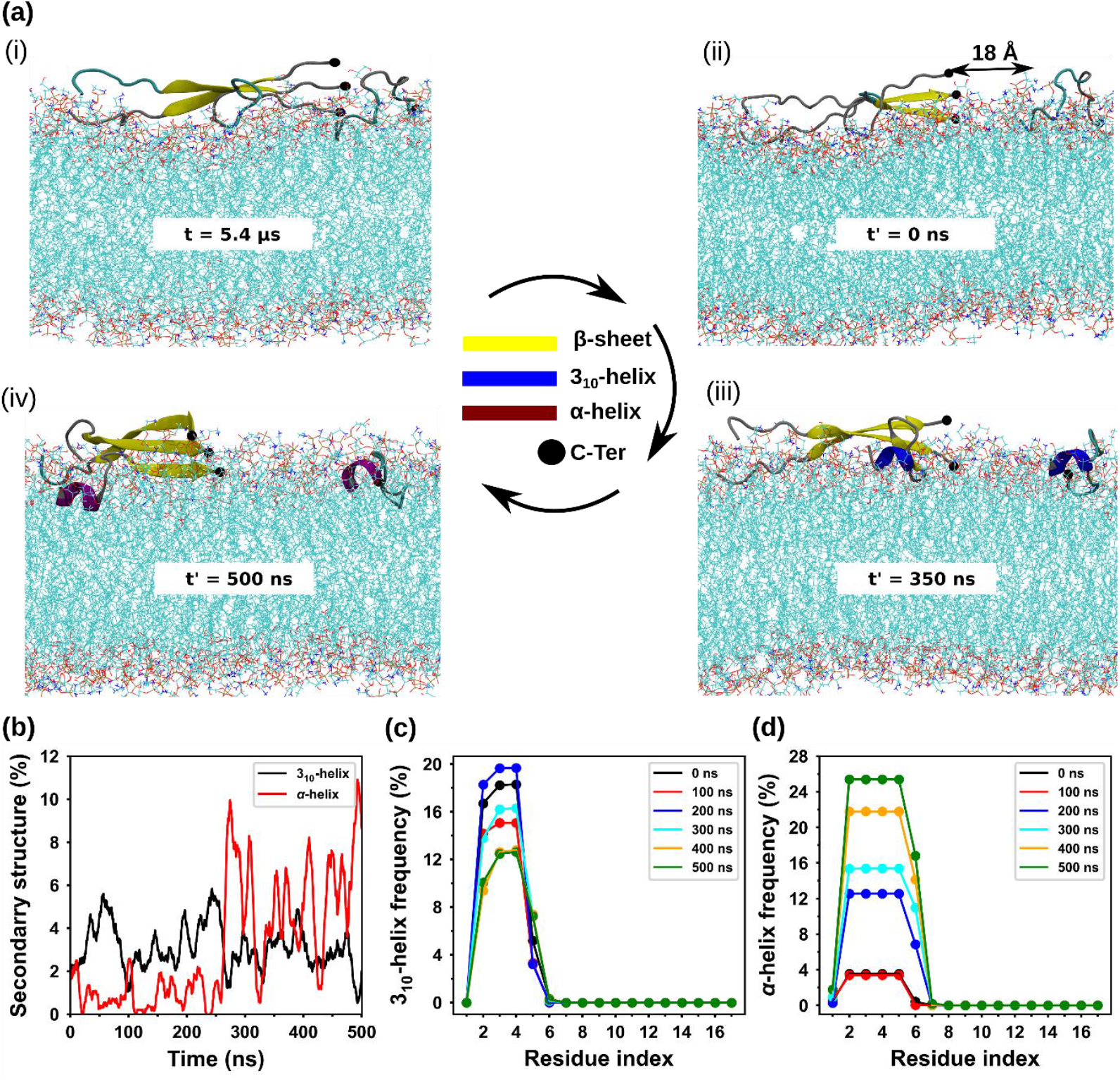
Snapshots highlighting the shifting of the dissociated U3.5-NH_2_ peptide and the subsequent conformational transitions of both monomer and remaining trimer after continuing the simulation. The initial distance between the monomer and trimer was 8 Å shown in (a-(i)), and after displacement of the monomeric peptide to a distance of 18 Å (a-(ii)), the evolution of a 3_10_-helix (a-(iii)) and α-helix (a-(iv)). Whereas, *t* represents time for the initial tetramer simulation in Figure 2, *t*^*′*^ represents time for the “re-started” simulation described here. (b) Re-starting the simulation for a further 500 ns, the secondary structure evolution is shown. (c) The 3_10_-helix and (d) α-helix frequency as a percentage for individual residues and at different times during the simulation.

Previously, we have shown that the coil-to-helix transition in U3.5-NH_2_ begins at the N-terminus, with 3_10_-helices acting as precursors in the formation of α-helices^5^. By the end of 500 ns, the single peptide contained 8–10% α-helix, suggesting that its interaction with the lipid bilayer induced helix formation, similar to the behaviour observed in the single peptide simulations (**Figure 1**). However, the dynamics of the transition was slower, likely due to the presence of the trimer in its vicinity. Unlike the U3.5-NH_2_ monomer in **Figure 1**, the single peptide was attached strongly to the bilayer before separating from the tetramer. In this case, the dynamics of the coil-to-helix transition would depend on the weakening of electrostatic interactions between the cationic U3.5-NH_2_ residues and the phosphate groups on the lipids. Notably, the trimer remained intact during the extended simulation, though intermittent formation of partial α-helix was observed (**Figure 3(a)**) suggesting that the trimeric moiety was beginning to lose a further peptide unit. It was reasonable to expect complete dissolution of the trimer at the bilayer surface over an extended period of time. Therefore, a simulation was designed with four *monomeric* peptides on the bilayer surface, each with partial helical content (end-state of the separated U3.5-NH_2_ peptide in **Figure 3(b)**).

#### Independent simulation of four Uperin 3.5-NH_2_ peptides at a membrane interface

Four U3.5-NH_2_ peptide monomers, with partial helical content corresponding to the final state in **Figure 3**, were randomly placed on the bilayer surface with an average inter-peptide separation of 30 Å (**Figure 4(a)**). Each monomer contained approximately 8-10% α-helix content. Figure 4(a) illustrates these peptides initially placed with partial helices on the lipid surface (top view of the membrane) as representative snapshots of the conformational changes of U3.5-NH2 peptides from 0-2800 ns simulation time. Initially, the peptides exhibited partial helicity, followed by a rapid increase in helical content at later stages of the simulation. Similar to the analysis used above, the secondary structure was quantified during the simulation, and the time-averaged frequencies of 3_10_-helix formation, calculated over 100 ns intervals, indicate dynamic transitions between 3_10_-helix and α-helix states over the first 1000 ns. After 1000 ns, the frequency of 3_10_-helix formation decreased, correlating with a stable transition into the α-helical structure across each residue for all four U3.5-NH_2_ peptides. The stabilised α-helical conformation exhibited a higher frequency, indicating the dominance of this secondary structure at the membrane surface in the later stages of the simulation (**Figure 4(b-d)** and **Figure S7**). Therefore, U3.5-NH_2_ peptides adopted an α-helical conformation at the lipid bilayer surface after dissociating from the initial amyloid state. These amphipathic α-helices lay flat at the bilayer surface with their hydrophobic sides facing the hydrophobic interior of the bilayer, shown in **Figure S8**, suggesting a carpet-like mechanism of anti-bacterial action. The series of simulations, from the partial dissolution of the amyloid in **Figure 2** to the helical transformations of U3.5-NH_2_ peptides in **Figures 3 and 4**, suggest a mechanism of antimicrobial action in which the amyloid state serves as a stable storage vehicle for antimicrobial peptides.

**Figure 4:**
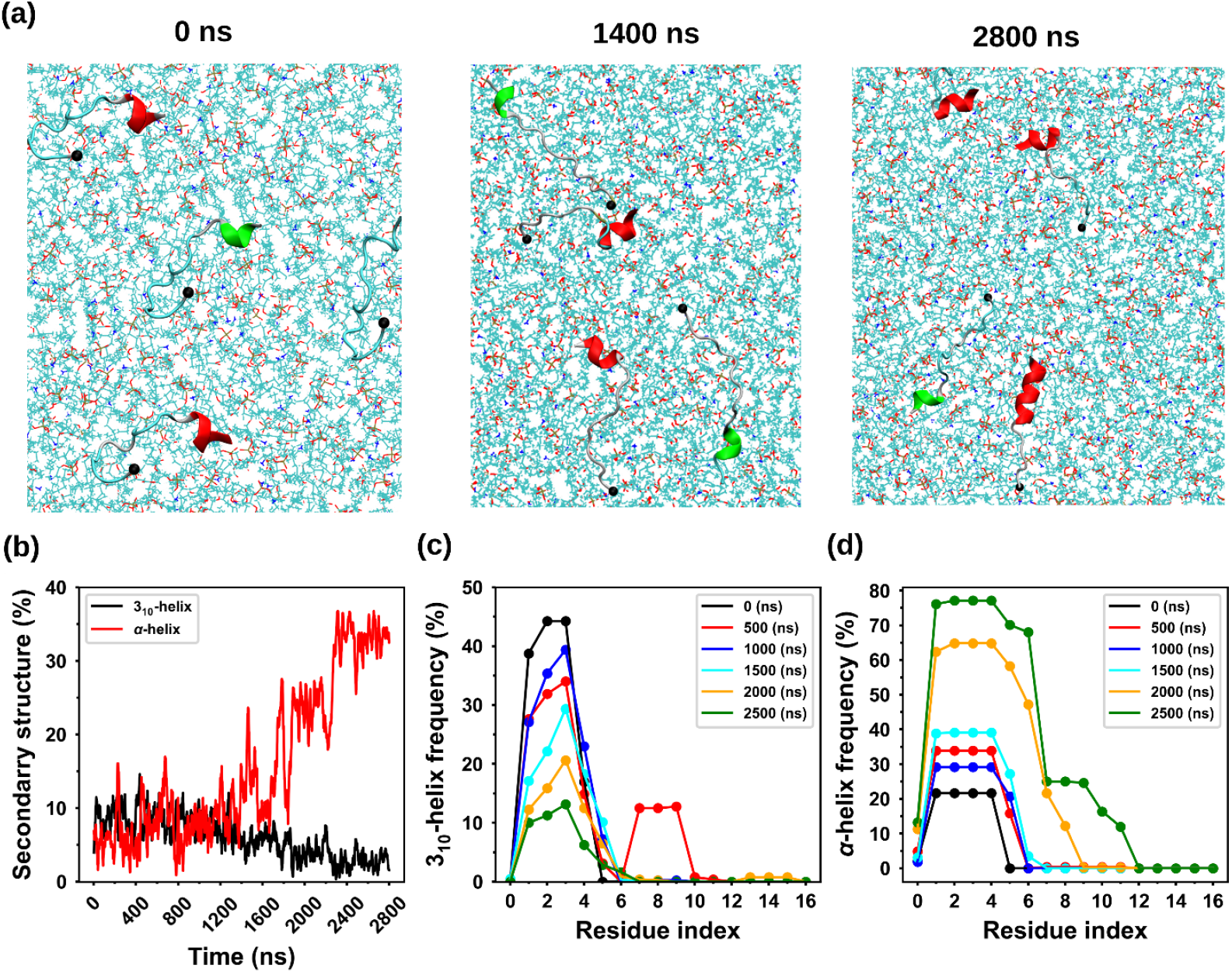
Structural evolution of U3.5-NH_2_ peptides after four U3.5-NH_2_ peptides were placed randomly at the surface of the lipid bilayer simulations, top view of the lipid surface. (a) Representative snapshots illustrate the conformational changes of U3.5-NH2 peptides. Initially, the peptides exhibit partial helicity, followed by a rapid increase in helical content at later stages of the simulation. (b) Secondary structure is quantified during the simulation and the time-averaged frequencies of 3_10_-helix formation, calculated over 100 ns intervals, indicate dynamic transitions between 3_10_-helix and α-helix states over the first 1000 ns. (c) After 1000 ns, the frequency of 3_10_-helix formation decreases, correlating with a stable transition to the α-helical structure across each residue in the U3.5-NH_2_ peptides. (d) The stabilized α-helical conformation exhibits higher frequency, indicating the dominance of this secondary structure at later stages of the simulation.

A Ramachandran plot corresponding to the four-peptide simulation in **Figure 4** is shown in **Figure S9**. The distribution of dihedral angles (Φ and Ψ) during the simulations revealed that the angles predominantly occupy allowed regions for α-helix and β-sheet secondary structure and reflecting the structural stability of the peptide conformations (**Figure S9(a)**). A potential of mean force (PMF) was calculated using the Boltzmann inversion of the distribution of Φ and Ψ angles from **Figure S9. Figure 5** illustrates the free energy landscape (PMF) corresponding to conformational transitions plotted with the dihedral angles, Φ and Ψ, as reaction coordinates. The free energy landscape revealed a global minimum basin corresponding to the right-handed α-helical region, which indicates the α-helical structure as the most stable conformer in the membrane environment. However, a second energy minimum was observed in the β-sheet region of the Ramachandran plot, which possibly appeared due to the extended peptide conformation anchored to the membrane surface. In addition, a PMF was also calculated using the 3_10_-helix and α-helix populations as reaction coordinates, (**Figure S9(b)**). This energy landscape highlighted the relative stabilities and facile transitions between the 3_10_-helix and α-helical conformations possible for the U3.5-NH_2_ peptides. Figure S10 shows the Ramachandran and PMF plots at the beginning and end of the 2.8 µs simulation, respectively. These calculations confirmed that α-helix is the most energetically favourable state for U3.5-NH_2_ peptides on the lipid bilayer. The peptide-membrane interactions also facilitated the structural transition of U3.5-NH_2_ peptide from a random coil to α-helical structure ^5^.

**Figure 5.**
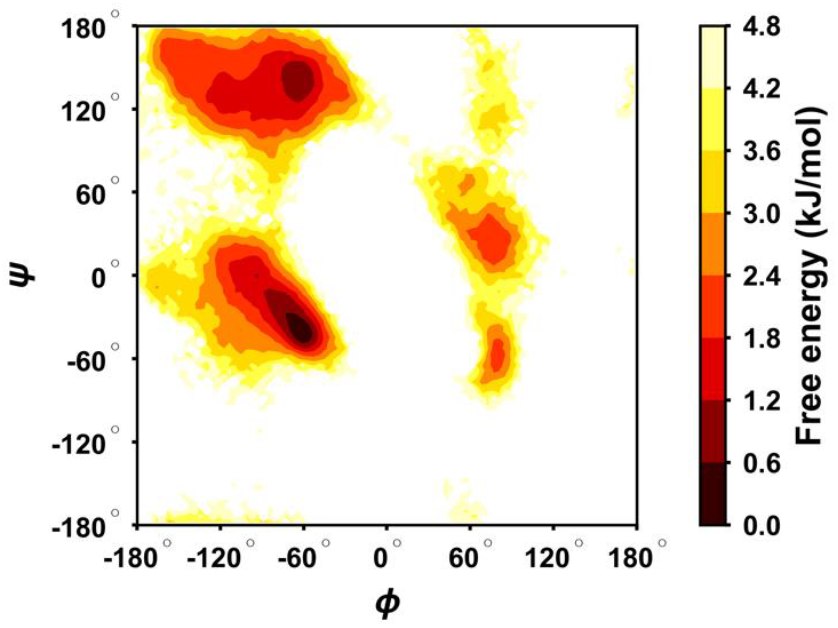
Free energy landscape of Φ and Ψ dihedral angles after four U3.5-NH_2_ peptides were placed randomly at the surface of the lipid bilayer simulations. The free energy landscape, derived from the Boltzmann distribution of Φ and Ψ angles, reveals a global minimum basin corresponding to the right-handed α-helical region. A second energy minimum is observed in the β-sheet region of the Ramachandran plot, representing a metastable state.

#### Uperin3.5-NH_2_ – lipid interactions

In addition to secondary structure transitions in the peptide, peptide-bilayer interactions also altered the organization and topology of the lipid bilayer. To further investigate these interactions, a lipid distribution analysis was performed for the 2.9 µs simulation in **Figure 4**. The number densities of the DOPG polar headgroups at the initial and final stages of the simulation (each averaged over a 100 ns interval) are shown as heat maps in **Figures 6(a), (b)**, respectively. At the final stage (**Figure 6(b)**), the redistribution of DOPG molecules was evident, with increased co-localization (association) with the peptide helices.^42^ The RDF plots (**Figures 6(c), (d)**) show the interaction between phosphates of DOPG and POPE with the peptide, highlighting a stronger correlation between DOPG molecules and peptides, particularly during their transition to an α-helical conformation in the later stages of the simulation.

**Figure 6:**
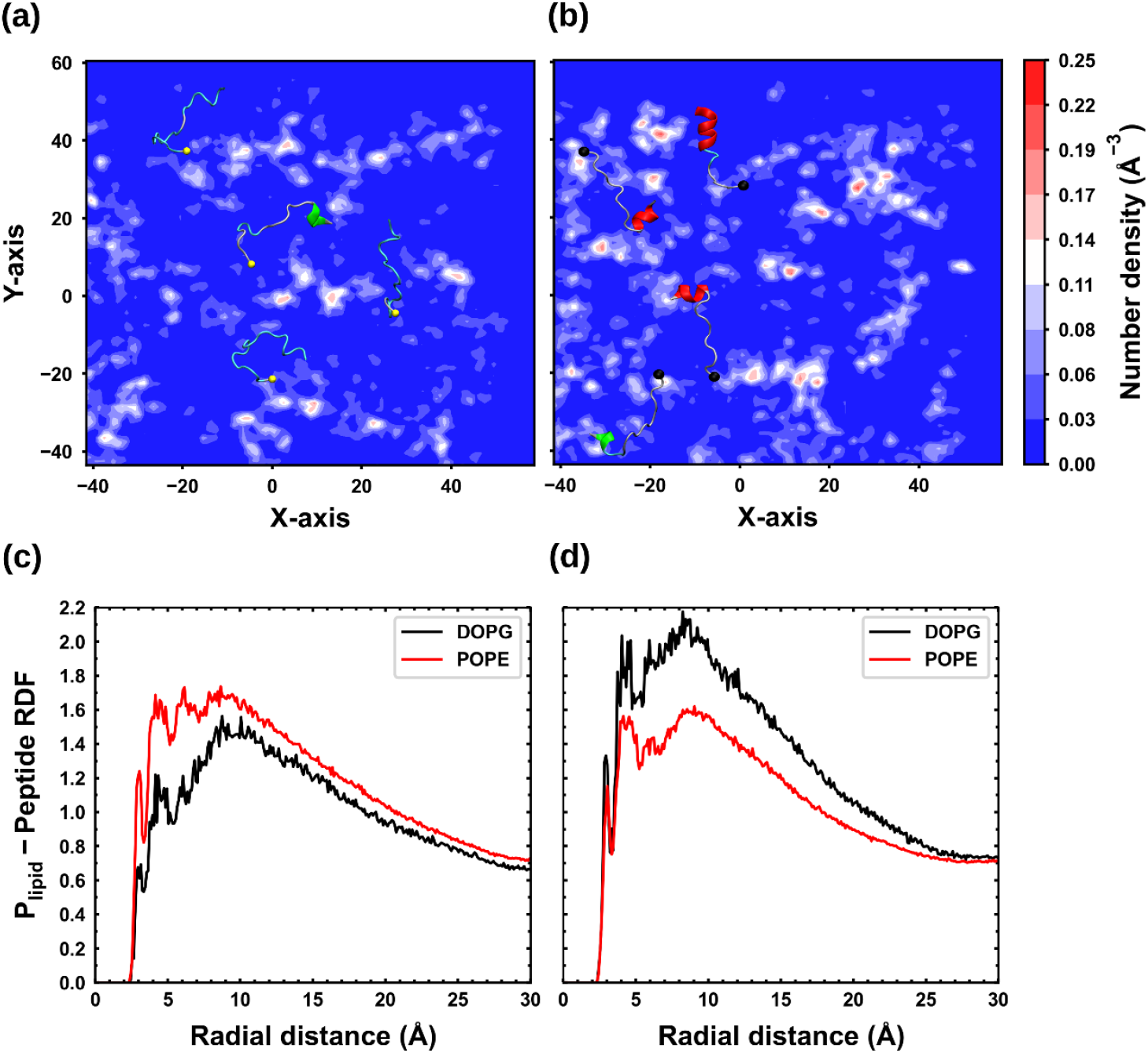
Lipid distribution analysis through number density and radial distribution function (RDF). (a) The number density of the polar headgroups of DOPG, averaged over 100 ns, is plotted for the initial and final stages of the simulation. At the final stage (b), the redistribution of DOPG molecules is evident, with increased association around the peptide helices. (c) The phosphate RDFs of DOPG and POPE highlight the enhanced correlation between DOPG molecules and the peptides, particularly during their transition to an α-helical conformation.

Both the density map in **Figure 6(b)** and the RDF in **Figure 6(d)** indicate enhanced DOPG-peptide interactions and their co-localization within the membrane. The structural transition of the peptides to α-helices significantly altered lipid distribution within the bilayer. Additionally, peptide-membrane interactions led to the embedding of peptides into the bilayer. Together, these effects – lipid redistribution and peptide incorporation into the bilayer – likely contributed to changes in bilayer topology. The mean curvature (*H*) of the bilayer’s upper and lower leaflets was estimated at the beginning and end of the simulation (averaged over 100 ns, **Figure 7**). Initially, both leaflets exhibited nearly uniform mean curvature values corresponding to a flat membrane. However, by the end of the simulation (2.8 µs), significant differences emerged between them. In the upper leaflet, regions containing helical peptides were characterized by large positive *H* values, indicating localized changes in membrane curvature. These changes in curvature may result in localized membrane thinning and ultimately lead to membrane rupture.

**Figure 7:**
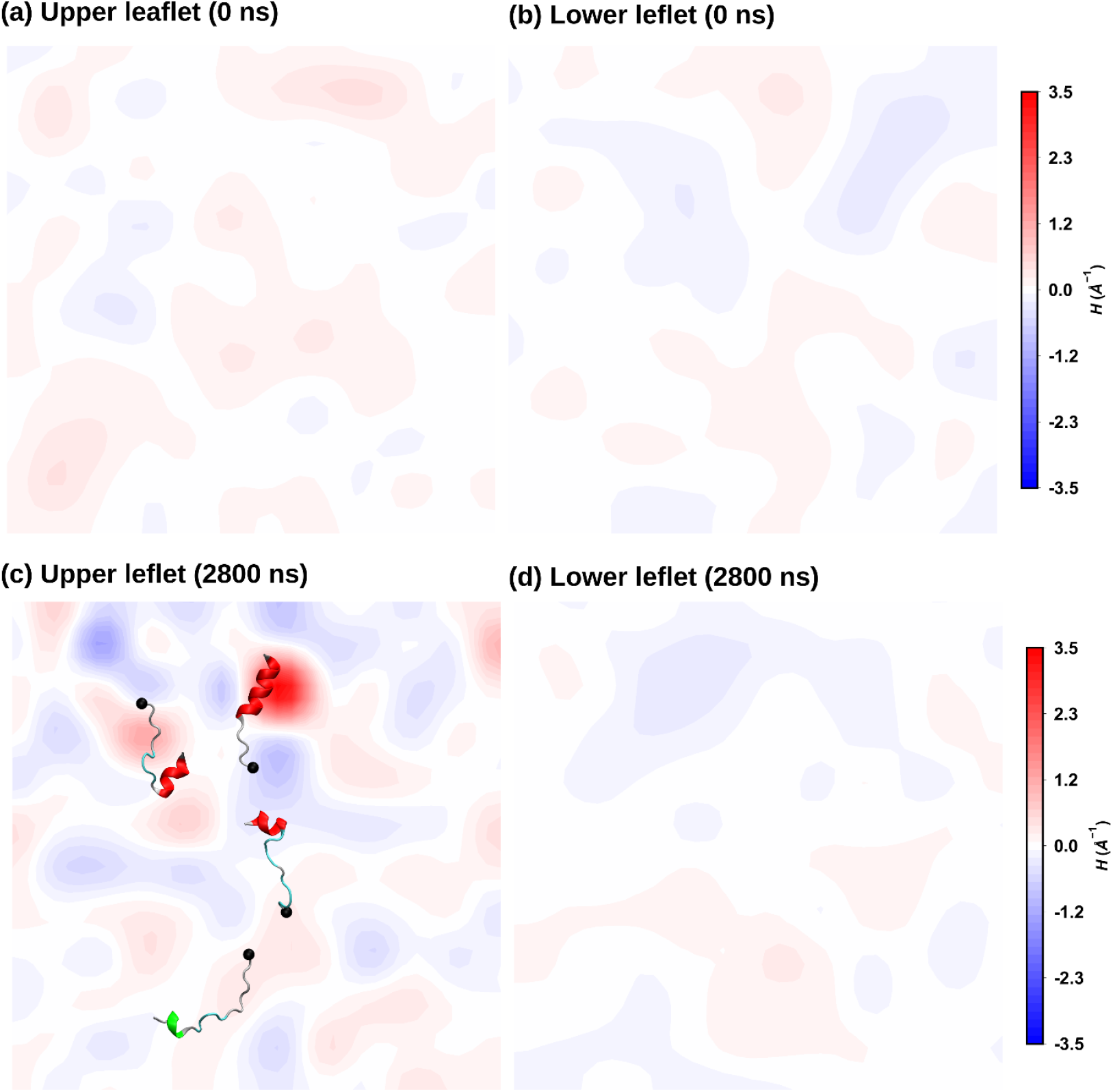
Mean curvature analysis of the lipid bilayer during peptide simulations. The mean curvature (***H***) was calculated at the initial and final stages of the simulations to assess lipid bilayer perturbations induced by the peptides. (a, b) At the initial stage, both leaflets exhibit low ***H***, indicating minimal impact on membrane curvature. (c) By the final stage (2800 ns), significant curvature alterations are observed in the upper leaflet, with pronounced positive and negative ***H*** values, highlighting the peptides’ disruptive effect on the bilayer structure. Note: The curvature has been plotted in the same scale, thus appears faded compared with the upper leaflet at the end of the simulation.

In summary, the monomer simulation showed that U3.5-NH_2_ rapidly attached to the lipid surface, forming a stable amphipathic α-helix that reoriented hydrophobic residues toward the bilayer interior and remained membrane-associated, while U3.5-OH exhibited only partial helical structures, fluctuating without clear insertion or stable secondary structure throughout the simulation. Furthermore, analysis of Cα atom distances revealed that U3.5-NH2 maintained strong and uniform membrane interactions (<5Å), indicating effective binding, while U3.5-OH exhibited weaker, fluctuating interactions (5Å–15Å), primarily via its N-terminus due to positively charged residues. The tetramer simulations revealed the β-sheet tetrameric U3.5-NH2 binds strongly to the bilayer, embedding its C-termini over time, while U3.5-OH remains weakly adsorbed with its C-termini oriented away from the bilayer. This suggests the C-terminal amidation enhances both peptide-bilayer and peptide-peptide interactions, stabilising β-sheet structure for a significant period of time.

The U3.5-NH2 tetramer began dissolving at ∼4.5 µs, with a single peptide separating but remaining nearby in a random coil state, contrasting with a single U3.5-NH2 peptide, which rapidly formed an α-helix upon bilayer adsorption; this suggests strong peptide-peptide interactions slow structural transitions, but peptide-lipid interactions ultimately drive tetramer dissociation and bilayer embedding. To enhance the conformational analysis of the dissociated peptide, the separated U3.5-NH2 peptide was repositioned, triggering a coil-to-helix transition over 500 ns, with 3_10_-helices forming first, followed by α-helices, while the remaining trimer stayed intact but showed signs of further dissociation, suggesting gradual peptide embedding into the bilayer. For comparison, an independent simulation of four Uperin 3.5-NH2 peptides on a membrane revealed a gradual transition from 3_10_-helix to stable α-helices, with α-helical conformations being the most energetically favourable state, supporting their carpet-like antimicrobial mechanism and the amyloid state as a peptide reservoir. Finally, Uperin 3.5-NH2 peptides altered lipid bilayer organisation by redistributing DOPG molecules, enhancing peptide-lipid interactions, embedding into the membrane, and inducing local curvature changes, which may contribute to membrane thinning and rupture.

## Conclusion

This study demonstrates that U3.5-NH2 peptides exhibit strong, stable interactions with lipid bilayers, rapidly adopting an amphipathic α-helical structure and maintaining membrane association, while U3.5-OH peptides show weaker, fluctuating interactions with only partial helical structures. The C-terminal amidation of U3.5-NH2 enhances both peptide-bilayer and peptide-peptide interactions, stabilizing its β-sheet structure and leading to gradual embedding into the bilayer during tetramer dissolution. Simulations further revealed that the dissociation of the tetramer triggers a coil-to-helix transition in the separated peptide, supporting a gradual embedding mechanism driven by peptide-lipid interactions. In comparison, independent simulations of four U3.5-NH2 peptides confirmed that their most energetically favourable state is the α-helix, which supports the antimicrobial “carpet-like” mechanism. Additionally, the peptides altered lipid bilayer organization by redistributing DOPG molecules and inducing local curvature changes, potentially contributing to membrane thinning and rupture. Overall, the study highlights the critical role of C-terminal amidation in stabilizing the peptide’s structure and promoting effective antimicrobial action.

This study suggests that amidation of the Uperin 3.5 peptide is crucial to its’ membrane interaction, hence implicated in the antimicrobial activity of the Uperin peptide. The amyloid state stabilises the Uperin peptides in the aggregated state and may prevent peptide degradation by mechanisms, such as proteolysis. Clearly, there is an impact of the higher charge in the aggregated state, which results in strong adsorption of the Uperin tetramer (amyloid) to the bilayer, although the non-amidated Uperin 3.5 orients, such that the significant negatively charge lies remotely form the membrane which impacts structural transitions that are observed for the amidated variant. Furthermore, this allows for the possibility of greater peptide-lipid interaction as the aggregate remains in contact with the bilayer surface. In a competition between peptide-peptide and peptide-membrane interactions, the latter dominate leading to gradual dissolution of the tetramer into individual peptides. These peptides undergo a coil-to-helix transition on the bilayer surface leading to formation of amphipathic peptide helices that partially embed themselves into the bilayer. These helices may lead to membrane disruption by either a carpet mechanism or detergent action. In terms of understanding the role of amidation versus non-amidation on the membrane activity of Uperin 3.5 peptide, our data suggest that the change in total peptide charge between amidated and non-amidated peptides is less important; illustrated by the simulation data for the aggregated tetramer, but that the amidation (albeit neutral) at the C-terminus of the peptide facilitates the necessary transition to an α-helix (via 3_10_-helices) with the membrane surface.

