## Supplementary material for "C-Terminal Amidation: Structural Insights into Enhanced Antimicrobial Peptide Efficacy and Amyloidogenesis": Supplemental Files.pdf

### Supplementary Information

#### Table of Contents


| | | Page no. |
|---|---|---|
| **Table S1** | Components included in MD Simulations of U3.5 near lipid bilayer. | S2 |
| **Table S2** | Initial parameters used for U3.5 monomer and tetrameric MD simulations. | S2 |
| **Figure S1** | Major components used for lipid bilayer-peptide interaction MD simulations. | S3 |
| **Figure S2** | Graphical representation for secondary structure evolution (α- and $3_{10}$ helices) of U3.5-$NH_2$ and U3.5-OH monomer simulations. | S4 |
| **Figure S3** | Average distance of the U3.5 peptide backbone ($C_\alpha$) amino acids from the membrane surface over the trajectory of monomer simulations. | S5 |
| **Figure S4** | Electrostatic potential map across the membrane bilayer during monomer simulations for U3.5-$NH_2$ and U3.5-OH. | S6 |
| **Figure S5** | The radial distribution functions (RDFs) compare phosphorus atoms (P) near the Cα atoms of specific residues. | S7 |
| **Figure S6** | Major contributions to Van der Waals (VdW) and electrostatic interaction energies between peptide-peptide and peptide-lipid during the simulation. | S8 |
| **Figure S7** | Secondary structure evolution of four U3.5-$NH_2$ peptides placed randomly at the surface of the lipid bilayer. | S9 |
| **Figure S8** | Snapshot at the end of the four-peptide simulation (2300 ns), showing the peptides are oriented with the hydrophobic surface within the lipid-bilayer. | S10 |
| **Figure S9** | Ramachandran plot used for analysis of dihedral angle distribution and free energy landscape in four peptide monomer simulations. | S11 |
| **Figure S10** | Ramachandran plots and PMF plots in the initial and final stages of the four-peptide simulation. | S12 |

**Table S1.** Components included in MD Simulations of U3.5 near lipid bilayer.

| Peptide-Lipid bilayer Simulation parameters | | | | | | | | |
|---|---|---|---|---|---|---|---|---|
| C-terminus modification | Uperin 3.5 Peptide (initial secondary structure) | Total no. of lipids (3:1) | | Total no. of atoms | No. of water molecules | No. of $Na^+$ ions | No. of $Cl^-$ ions | Time (ns) |
| | | POPE | DOPG | | | | | |
| Amidated | Random coil | 252 | 84 | 112830 | 23283 | 138 | 66 | 500 |
| | Parallel tetramer | 252 | 84 | 120780 | 25658 | 141 | 69 | 500 |
| | Parallel dimer | 252 | 84 | 121011 | 25915 | 154 | 76 | 500 |
| Non-amidated | Random coil | 252 | 84 | 112742 | 23254 | 141 | 59 | 500 |
| | Parallel tetramer | 252 | 84 | 120830 | 25676 | 142 | 66 | 500 |
| | Parallel dimer | 252 | 84 | 121126 | 25954 | 156 | 76 | 500 |

**Table S2.** Initial parameters used for U3.5 monomer and tetrameric MD simulations.

| C-terminal modification | Simulation setup | Box Size ($Å^3$) | Peptide distance from lipid (Å) | NaCl Conc. (M) |
|---|---|---|---|---|
| **Amidated** | Random coil | 101×101×114 | ~15 | 0.15 |
| | Parallel tetramer | 101×101×124 | ~20 | 0.15 |
| | Parallel dimer | 101×101×132 | ~20 | 0.15 |
| **Non-amidated** | Random coil | 101×101×114 | ~15 | 0.15 |
| | Parallel tetramer | 101×101×124 | ~20 | 0.15 |
| | Parallel dimer | 101×101×132 | ~20 | 0.15 |

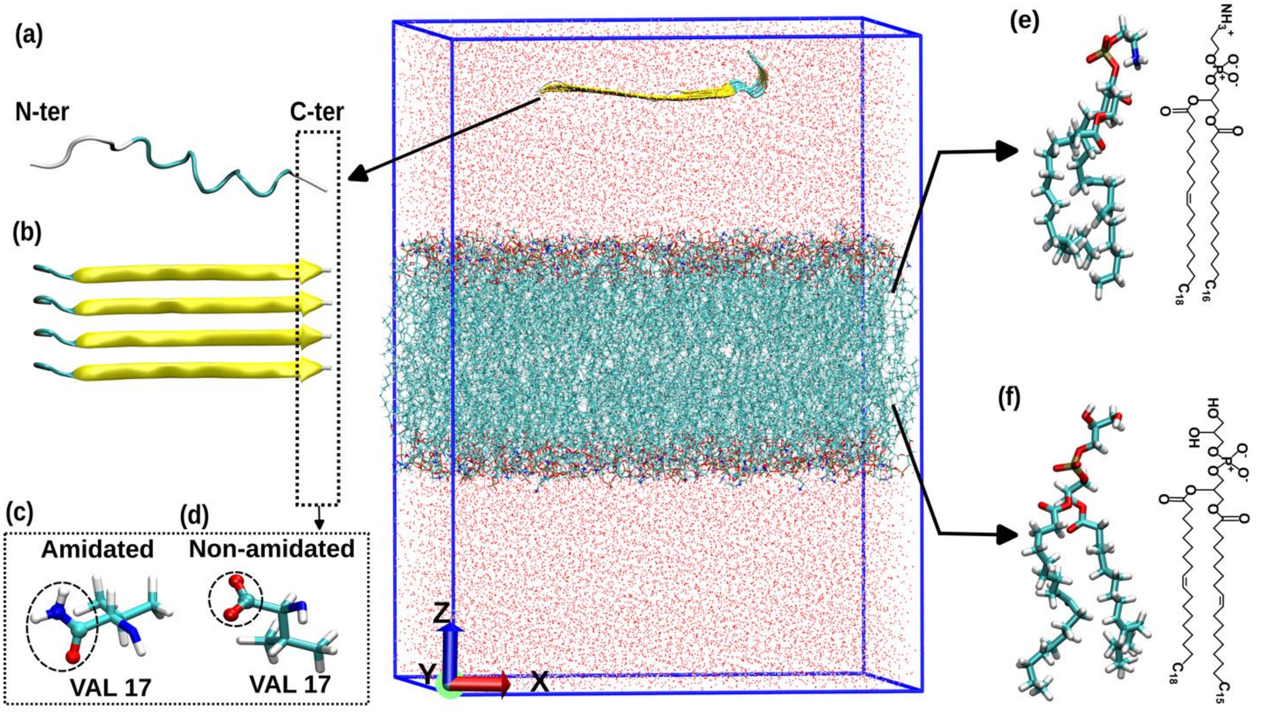


**Figure S1:** Major components used for lipid bilayer-peptide interaction MD simulations; **(a)** monomer peptide, **(b)** tetramer peptide, and **(c)** modification at the C-terminus at valine 17 (VAL17) shown by a dashed circle. The simulation box, as depicted in **(d),** contains the lipid bilayer with either a single peptide or a tetramer peptide from (c), as indicated in (a) and (b). The anionic lipid bilayer consists of POPE:DOPG 3:1, with their molecular structures shown in **(e),** and **(f)**, respectively.

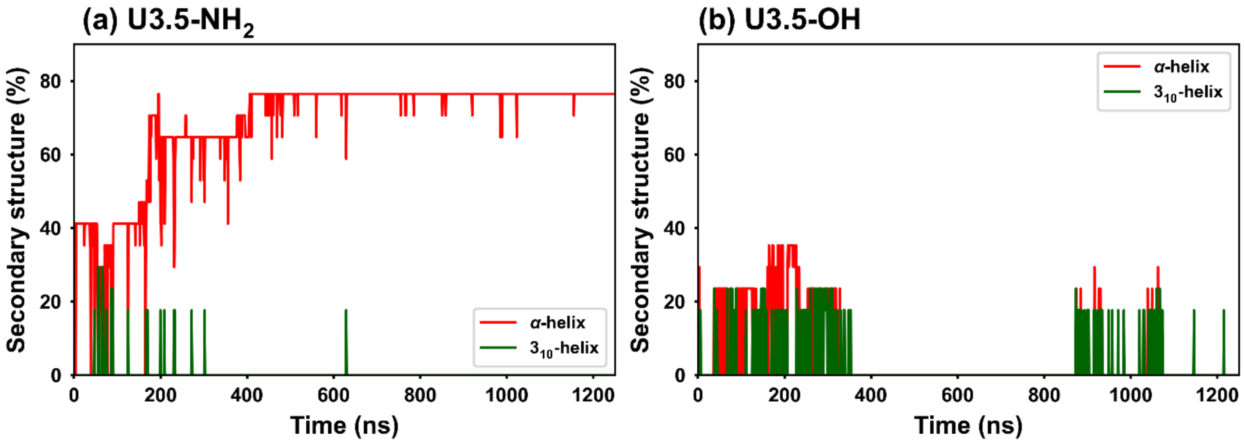


**Figure S2:** Secondary structure evolution during the MD simulation of Uperin peptide monomers at a lipid bilayer. (a) The U3.5-$NH_2$ peptide exhibits the transient formation of $3_{10}$-helix (green) and α-helix (red) during the initial stages of the simulation, eventually forming a stable, predominantly α-helical structure (~450-1200 ns). (b) In contrast, the U3.5-OH peptide fluctuates transiently between $3_{10}$-helix and α-helix structures throughout the simulation but fails to form any stable α-helical structures.

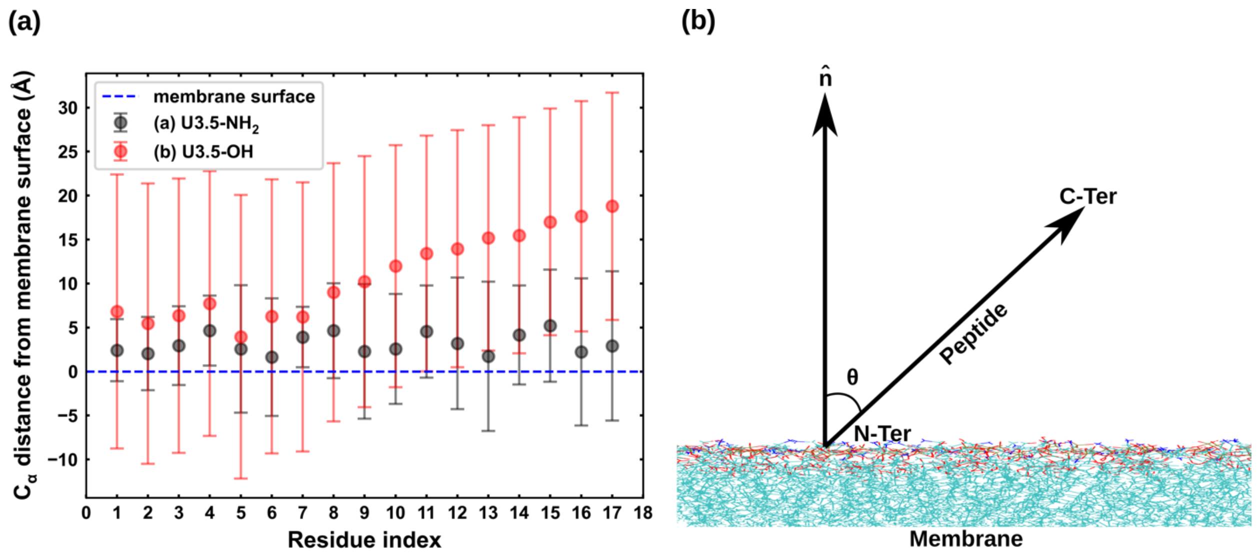


**Figure S3:** (a) Average distance of the U3.5 peptide backbone ($C_\alpha$) amino acids from the membrane surface over the trajectory of monomer simulations. The averaged Cα distance of the U3.5-$NH_2$ peptide indicates strong and consistent interaction across the entire peptide length, signifying effective membrane association. In contrast, the U3.5-OH peptide predominantly interacts through its N-terminus, particularly up to $C_\alpha8$, due to the presence of positively charged residues (ARG7 and LYS8) located towards the N-terminus. In the U3.5-OH case, the negatively charged C-terminus is repelled from the membrane, resulting in limited interaction. Moreover, the larger deviations from the mean distance observed in U3.5-OH highlights the weaker membrane interaction compared to U3.5-$NH_2$. (b) Graphic representation of the peptide orientation by the angle (θ) relative to the membrane surface.

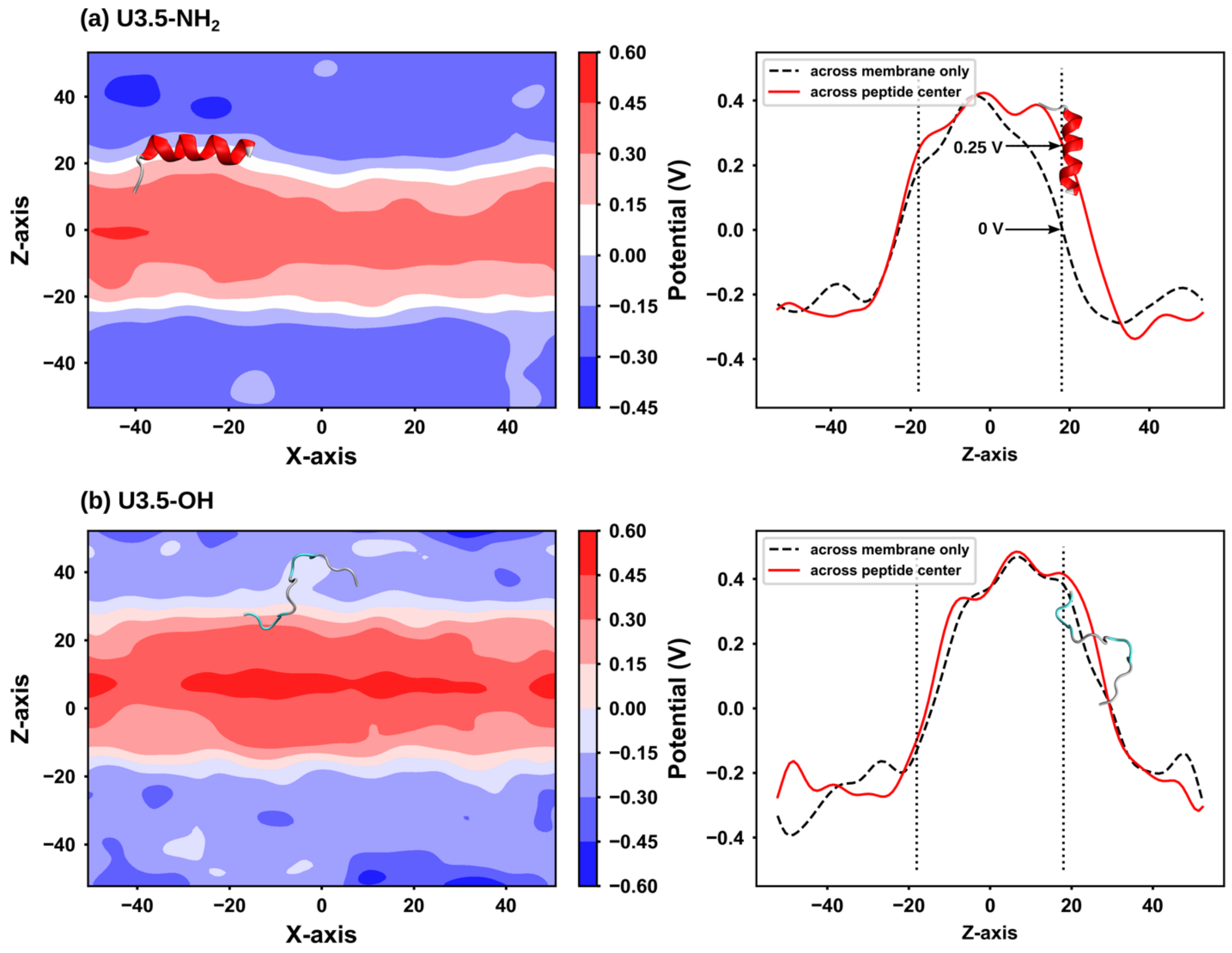


**Figure S4:** Electrostatic potential map across the membrane bilayer during monomer peptide simulations. (a) The electrostatic potential map of the U3.5-$NH_2$ peptide in the X-Z plane reveals a distinct positive region around X, Z = (−30, 20), corresponding to the peptide's location. Additionally, the potential profile along the Z-axis shows a clear difference in potential between the peptide centre and the membrane, particularly near Z = 20. (b) In contrast, the potential map for the U3.5-OH peptide-membrane system lacks a clear distinction in potential for the peptide centre and the membrane location. This observation suggests a dilution of the electrostatic potential, likely caused by the C-terminus of the U3.5-OH peptide well above the membrane surface, thereby leading to weaker interaction with the lipid bilayer.

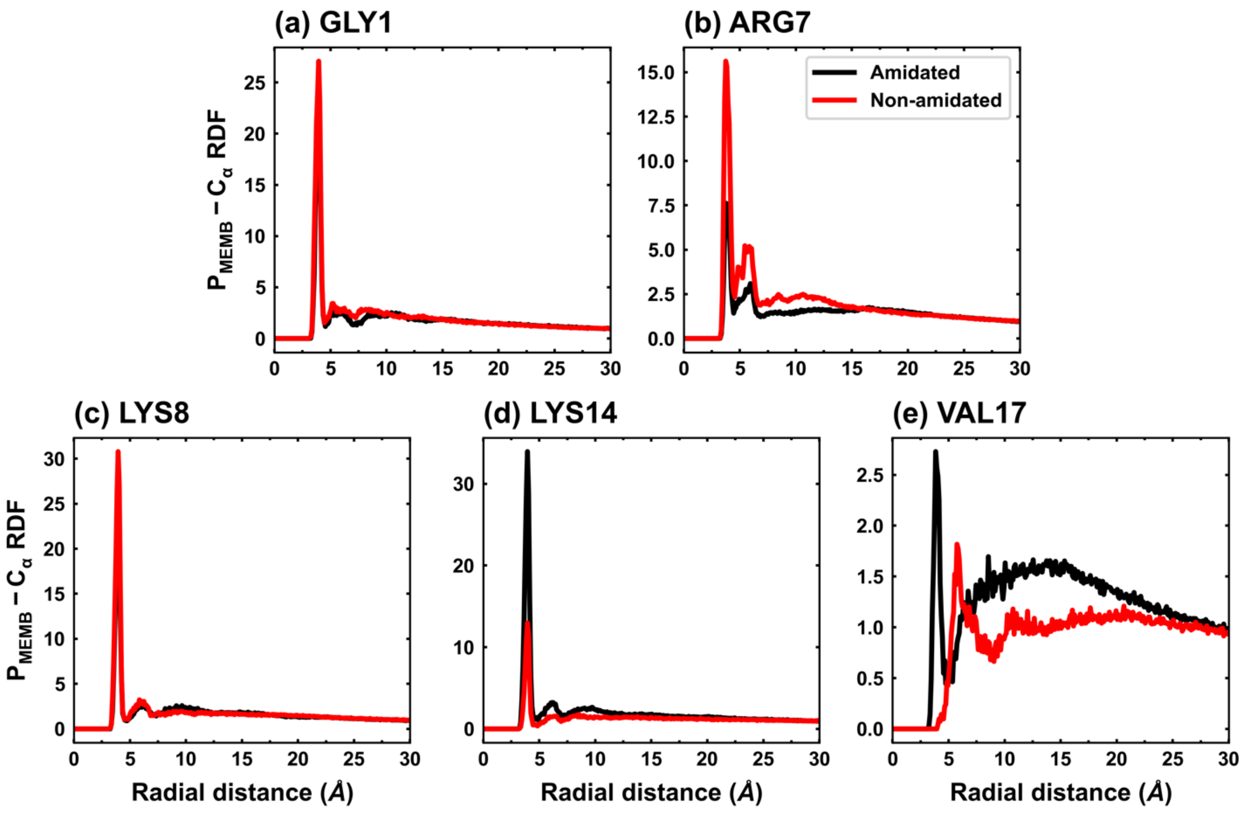


**Figure S5:** Distribution of lipid molecules around charged residues (including N- and C-termini) to assess the magnitude of peptide-lipid bilayer interactions. The radial distribution functions (RDFs) compare phosphorus atoms (P) near the Cα atoms of specific residues. (a, b) RDFs for positively charged residues (e.g., GLY1, ARG7 and LYS8) indicate a strong association with phosphate groups for both amidated ($U3.5\text{-}NH_2$) and non-amidated (U3.5-OH) peptides. (c, d) However, distinct differences are observed at the C-terminus, particularly for LYS14 and VAL17. The negatively charged C-terminus of the U3.5-OH peptide weakens its interaction with phosphate groups, highlighting reduced association strength compared to the amidated counterpart.

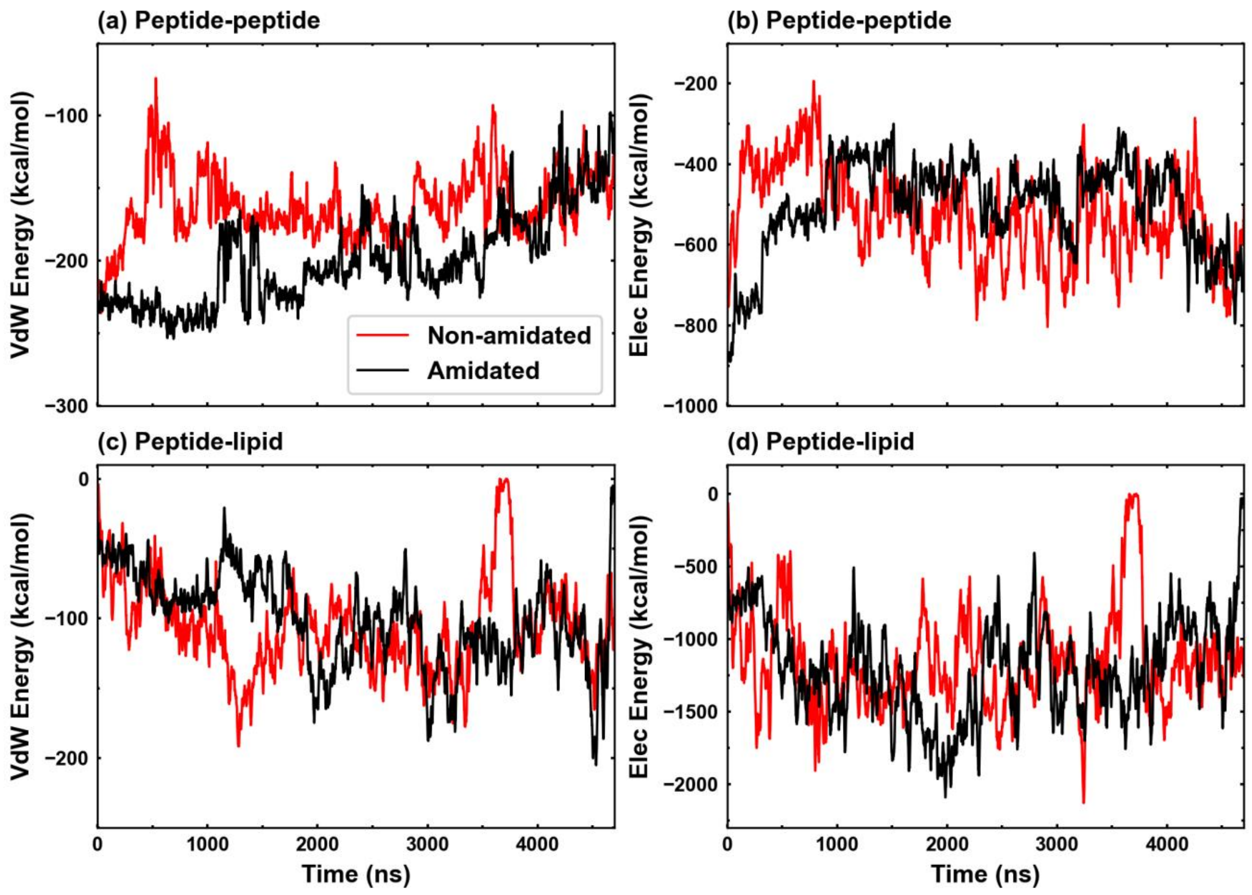


**Figure S6**: Major contributions to Van der Waals (VdW) and electrostatic interaction energies between peptide-peptide and peptide-lipid during the simulation. (a, b) VdW and electrostatic energy plots for peptide-peptide interactions of U3.5-$NH_2$ and U3.5-OH, respectively. (c, d) VdW and electrostatic energy plots for peptide-lipid interactions of U3.5-$NH_2$ and U3.5-OH, respectively.

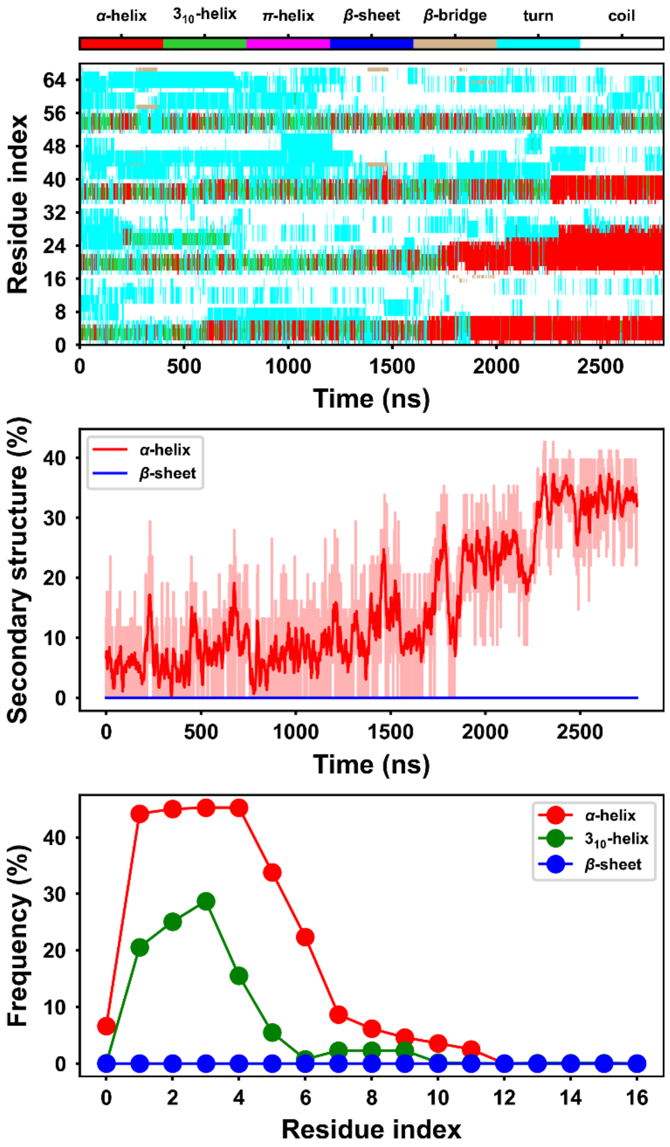


**Figure S7:** Secondary structure evolution of four U3.5-$NH_2$ peptides placed randomly at the surface of the lipid bilayer. The initial structure obtained from the peptide that dissociated from the tetramer in the β-sheet tetramer simulation. (a) The secondary structure heatmap illustrates the dynamic interconversion between $3_{10}$-helix and α-helix of each peptide (plotted as 0-16, 17-33, 34-50, 51-67 on y axis) over the first 1500 ns of simulation. Beyond this period, three of the four peptides where in an α-helical conformation until the end of the simulation. (b) The percentage of α-helix content shows a gradual increase during the later stages of the simulation, reflecting the stabilization of $3_{10}$-helix into the α-helical structure. (c) The frequency analysis of secondary structures indicates a prominent role of the peptide's N-terminus in driving the secondary structure evolution.

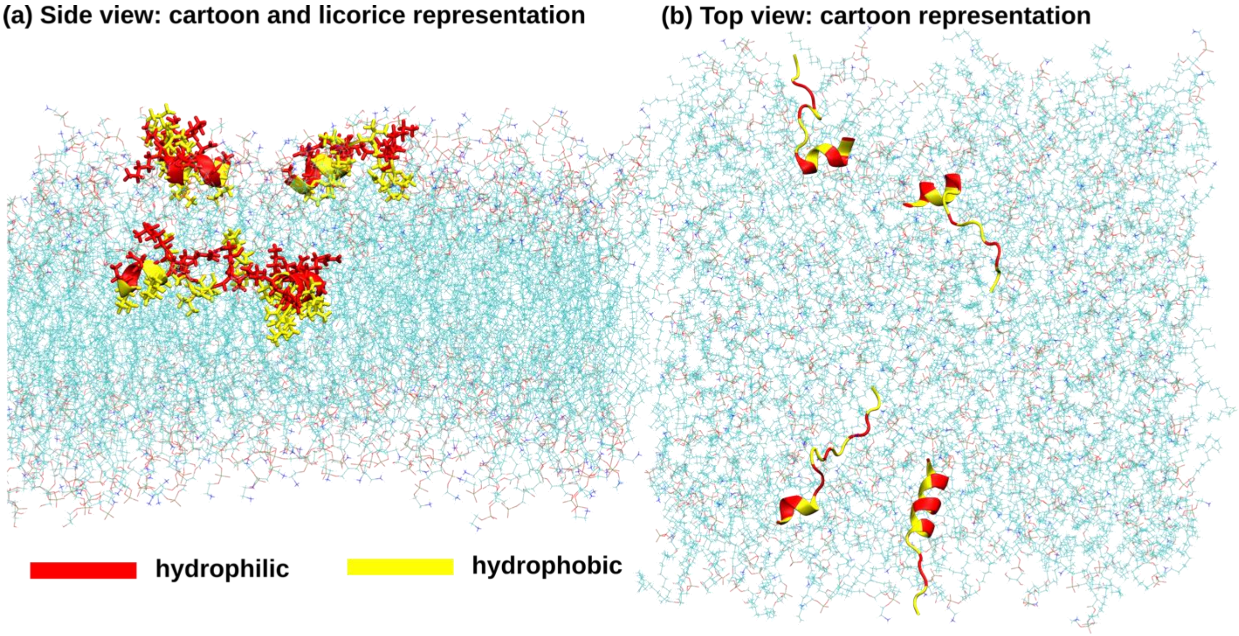


**Figure S8:** Snapshot at the end of the four-peptide simulation (2300 ns), showing the peptides are oriented with the hydrophobic surface within the lipid-bilayer, showing the (a) side-view pf the lipid bilayer and (b) top-view.

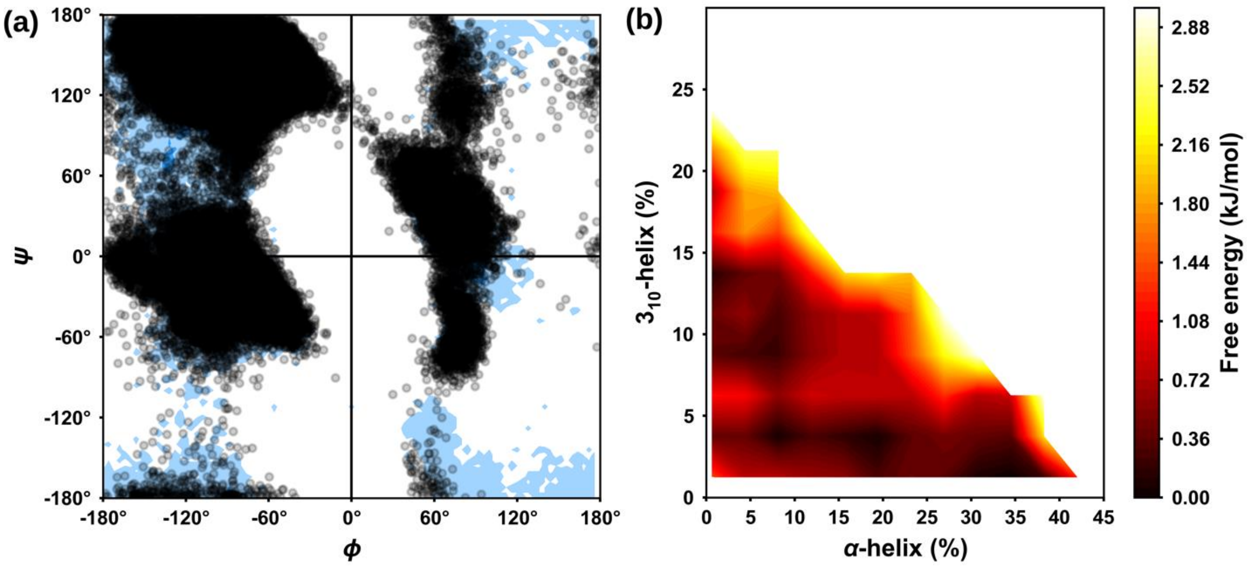


**Figure S9:** Ramachandran plot used for analysis of dihedral angle distribution and free energy landscape in four peptide monomer simulations. (a) The distribution of dihedral angles ($\Phi$ and $\Psi$) during the simulations reveals that the angles predominantly occupy allowed regions for $\alpha$-helix and $\beta$-sheet secondary structure and reflecting the structural stability of the peptide conformations. (b) Using the $3_{10}$-helix and $\alpha$-helix populations as reaction coordinates, the free energy landscape was constructed based on the Boltzmann distribution. This landscape highlights the relative stabilities and transitions between the $3_{10}$-helix and $\alpha$-helical conformations.

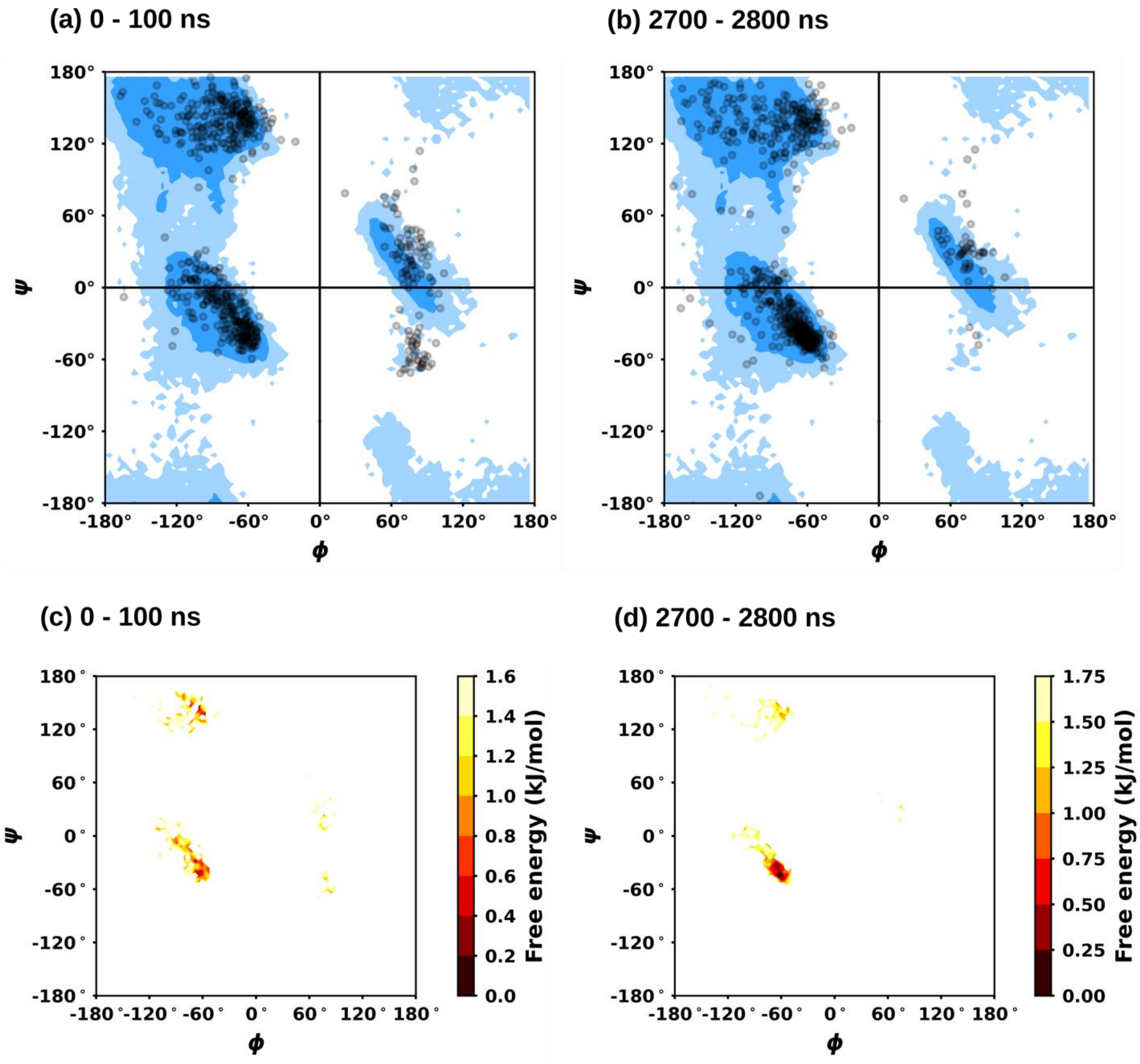


**Figure S10:** Ramachandran plots for the (a) initial 100 ns and (b) final 100 ns, in the four peptide-membrane simulations, and the corresponding potential of mean force (PMF) plots in (c) initial 100 ns and (d) final 100 ns, respectively.
